# Tracking GAD-specific T-cell expansions in Type 1 diabetes by intradermal GAD-Alum challenge

**DOI:** 10.64898/2025.12.02.690944

**Authors:** Stephanie J. Hanna, Emma JS Robinson, Terri C Thayer, Maki Nakayama, Laurie Landry, Robert Andrews, Garry Dolton, Joanne Davies, Evangelia Williams, James A Pearson, Andrew K. Sewell, Parth Narendran, David Wraith, Alexandra Howell, Philippa Young, Mary Hart, Anton Lindqvist, F. Susan Wong, Tim IM Tree, Colin M Dayan, Danijela Tatovic

## Abstract

Identifying and monitoring autoreactive T cells that drive beta cell destruction remains a major obstacle to developing effective immunotherapies for type 1 diabetes (T1D). These cells are extremely rare in peripheral blood and cannot be accessed directly from the pancreas. We used intradermal injection of GAD-Alum to recruit GAD-specific T cells to accessible sites in the skin and skin-draining lymph nodes (LNs), sampled by skin suction blisters and ultrasound-guided LN aspiration. Peripheral blood samples obtained before GAD injection were restimulated with GAD *in vitro* to detect reactive CD4+ T cells. Re-expression of selected T cell receptors (TCRs) confirmed antigen specificity. Up to 70% of T cells at the skin injection site were clonally-expanded and 4 of 14 (28%) re-expressed TCRs were GAD-reactive. In draining LNs 1 of 14 (4%) clonally-expanded TCRs was GAD-reactive, representing ∼0.08% of all T-cells. GAD-reactive cells across compartments displayed Th1 and Th17-associated transcription signatures. These results demonstrate the intradermal autoantigen challenge, coupled with scRNAseq, enables direct identification and molecular profiling of autoreactive T cells *in vivo*. This minimally invasive approach provides a powerful platform for tracking antigen-specific specific T cells to monitor disease activity and evaluate immune interventions in T1D.

## Introduction

The licensing of Teplizumab to delay the onset of type 1 diabetes (T1D) has demonstrated the potential of immune therapies to postpone insulin dependence(1). To accelerate the development of disease-modifying therapies, it is essential to develop techniques that can monitor how interventions affect the pathogenic T cell response that drives beta-cell destruction. However, tracking antigen-specific T cells in T1D is extremely challenging due to their low frequency in peripheral blood (∼1:10,000)(2–4) and the inaccessibility of the pancreas and its draining lymph nodes in living donors. Several approaches have been used to overcome this limitation, including isolation of tetramer-positive cells from the peripheral blood(5, 6), *in vitro* activation assays to enrich antigen-specific cells(7) and TCR sequencing of islet infiltrating lymphocytes from deceased donors followed by re-expression in cell-based systems to confirm specificity(8). Each has significant drawbacks. Tetramer-based detection is limited by low frequencies, some non-specific binding and the need to predefine epitopes and HLA-restriction. ELISPOTs restrict phenotypic analysis to cytokine secreting cells, and prolonged *in vitro* culture can bias clone selection and distort the *in vivo* phenotype. Flow and mass cytometry have provided important insights into cell phenotypes in T1D(9, 10), but the limited panel sizes restrict discovery of novel biomarkers. There is therefore a clear need for methods that combine confident identification of disease-relevant antigen-specific T-cells with complete, unbiased phenotypic profiling that faithfully reflects their *in vivo* status. scRNAseq is ideally suited to this with early studies identifying predictive biomarkers in individuals who progress to T1D(11) and defining transcriptional pathways specific to islet-antigen reactive lymphocytes(7).

Developing approaches that allow reliable profiling of antigen-specific T cells in T1D will be key to monitoring and optimising how immunotherapies modify disease activity, identifying at-risk individuals and matching them to the most appropriate therapeutic strategy(12–15).

We recently reported that intradermal injection of a proinsulin peptide (C19-A3) conjugated to gold nanoparticles (GNP) induced marked clonal expansion of T cells at the injection site, with over 50% of expanded clonotypes specific for either proinsulin or the GNP core particle(16). The frequency was around 1000-fold higher than that seen in the peripheral blood. We also showed that intradermal injection of recall antigens such as PPD led to accumulation of antigen-specific cells in skin-draining lymph nodes within 3-5 days(17, 18). These findings led us to hypothesise that diabetes autoantigen-specific T cells could be recruited to accessible sampling sites by intradermal injection of whole autoantigens, enabling their isolation and characterisation *in vivo*(18).

Glutamic acid decarboxylase 65 (GAD) is a major autoantigen targeted by both T cells and B cells in T1D(19). Here, we combine scRNAseq with intradermal injection of GAD-Alum to identify and profile GAD-specific T cells in people with T1D using three complementary enrichment strategies: 1. Recruitment to the injection site, 2. Enrichment in the skin draining lymph node, and 3. *Ex vivo* activation and isolation of GAD-reactive peripheral blood CD4+ T cells.

## Results

### Response to intradermal GAD-Alum injection

All participants who received intradermal GAD-Alum developed an immediate local skin reaction characterized by a circular erythematous area approximately 2-3cm in diameter. The erythema resolved within 45-60 minutes under observation. A delayed asymptomatic erythematous skin induration of similar size then appeared at the injection site two days later and resolved completely within 15 days (Supplementary Figure S3).

### Response to intradermal GAD-Alum in the skin and skin-draining lymph nodes

We aimed to enrich for GAD-specific cells and be able to capture their *in vivo* phenotype without *ex avivo* expansion. Based on our previous work, we hypothesized that intra-dermal injection of GAD-Alum would recruit clonally-expanded T cells to the skin and skin draining lymph nodes (LNs).

Skin blisters were obtained from the injection sites of three participants with T1D (Donors C, D and and E), 5-13 days after GAD Alum administration. scRNAseq-derived TCR sequences were obtained from all three donors and gene expression (GEX) data were generated from blisters from Donors D and E (Fig 1A). LN aspirates were collected before and after GAD-Alum injection from five donors. We refer to these samples as LN_Pre and LN_Post. All blisters with GEX information demonstrated T cell infiltrations (Fig 1A, Supplementary Fig S4, S5). LNs contained abundant CD4+ T cells, smaller CD8+ T cell proportions and substantial B cell fractions. (Fig 1A, Supplementary Fig S5). Clonally-expanded T cells, defined by identical paired TCRα and TCRβ sequences, were mapped onto the UMAP plots (Fig 1B).

**Fig 1:**
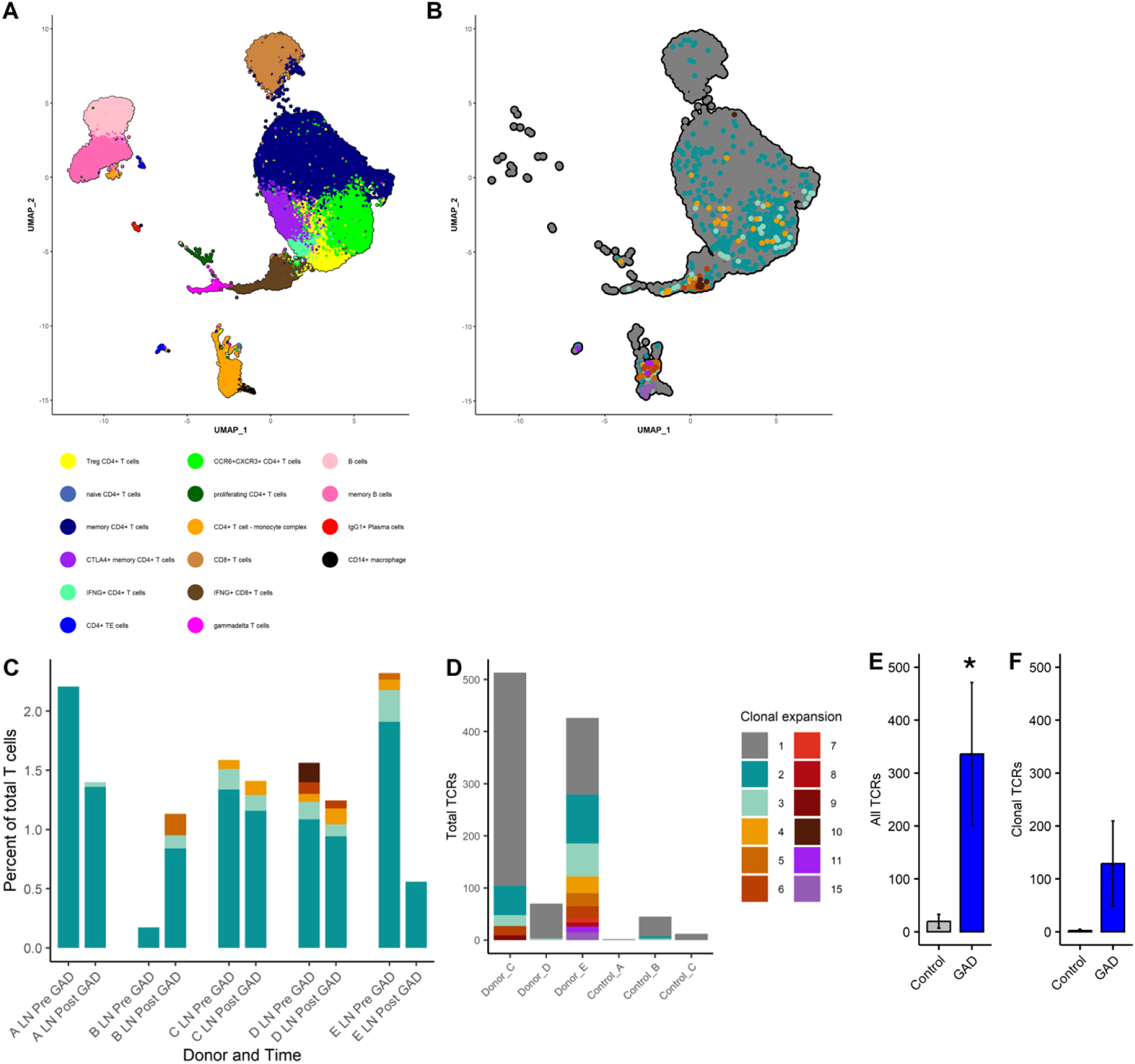
Single-cell RNA sequencing of lymph node and skin blister samples following intradermal GAD-Alum injection. **A**. UMAP plot showing all pre- and post-GAD lymph node (LN) cells and post-GAD blister cells, clustered in Seurat and coloured by cell type. **B**. UMAP plot of T cells expressing paired TCRs, coloured by clonal expansion number. **C**. Bar graph showing the proportion of clonally-expanded T cells in the LNs, expressed as a percentage of all T cells with sequenced TCRs and coloured by expansion size. D.Total number of TCRs sequenced from each of the skin blisters, coloured by clonal expansion number. (Clonal expansion legend colours refer to B-D) E. Total number of TCRs in blisters. F. Number of clonal TCRs in blisters.

Consistent with our previous findings(18, 20) we expected GAD-specific T cells to be enriched among clonally-expanded populations in the LNs. Clonal expansions were present both before and after intradermal GAD-Alum injection representing approximately 0.1-2.5% of all T cells with no significant change in abundance [Fig 1C,1D; Supplementary Fig 6]. Shared TCRs were largely restricted to samples from the same donor pre- and post-injection indicating persistence of certain clones between timepoints (data not shown). Donor D exhibited particularly high overlap between pre- and post-injection samples, coupled with the largest baseline clonal expansions (60 shared clonotypes across 156 cells from 14,708 total; Morista-Horn index 0.016; Fig 1C).

Donors C, D, and E all demonstrated substantial clonal expansions in skin blisters (∼20%, 5%, 45% of all T cells, respectively, exceeding those observed in LNs) (Fig 1D). Donor E also showed some evidence of antigen-presenting cells clustering with T cells, possibly reflecting ongoing physical interactions and antigen recognition *in situ*(21)).

Control blisters were collected from three donors with T1D who did not receive GAD-Alum. The total number of TCR-bearing cells was significantly higher in GAD-injected compared with control blisters (p<0.05, one-tailed t test), although the increase in clonally-expanded TCRs did not reach statistical significance (Fig 1E,F). The total number of cells recovered from the GAD-injected blisters was also increased compared to controls (data not shown)

### Re-expression of TCRs to confirm GAD specificity

The large clonal expansions observed in the skin strongly suggested an antigen specific response(18), which we sought to confirm by re-expressing the TCRs in a cell-based system. We selected TCRs with paired α and β chains representing the highest clonal expansion, including both CD4+ and CD8+ clonotypes where possible (Table 1). TCR transductants were first screened for activation by GAD-Alum presented by autologous LCLs and those showing a significant increase in NFAT reporter signal (data not shown) were subsequently tested against GAD-expressing LCLs. This analysis demonstrated that 4 of 14 (29%) of the blister-derived TCRs were GAD-specific, including at least one GAD-reactive TCR from each donor as indicated by increased fluorescence in the NFAT reporter assay (Fig 2A, Table1).

**Fig 2:**
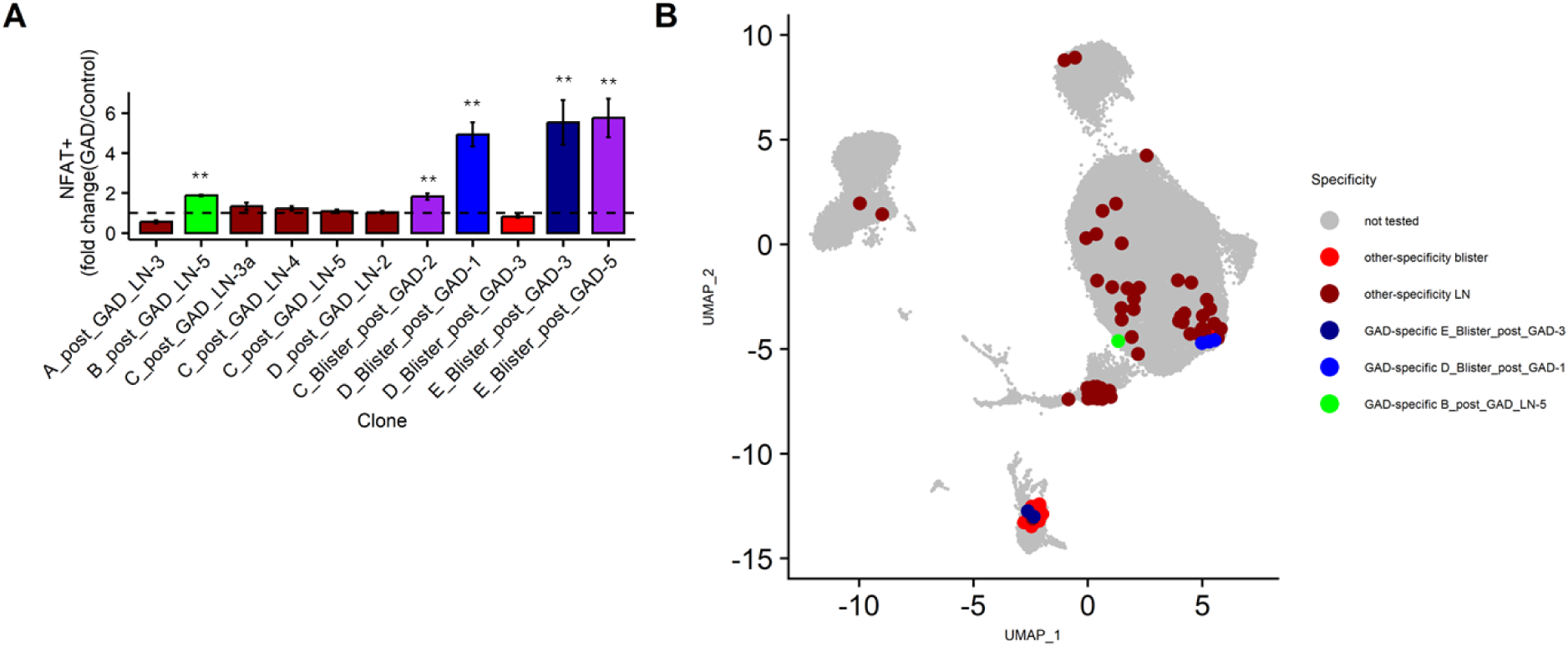
GAD-specificity of re-expressed TCRs. **A**. TCRs identified by scRNAseq were re-expressed in a 5KC reporter cells. TCRs showing positive responses when screened with GAD-Alum were re-tested using autologous GAD-expressing LCL lines. **p<0.01 one-tailed T test. Data are pooled from duplicate observations in three independent experiments. B UMAP plot showing the distribution of all re-expressed TCRs mapped back onto the single-cell dataset,illustrating phenotypes of GAD-specific T cells compared with those of other specificities, coloured by clone ID.

**Table 1:**
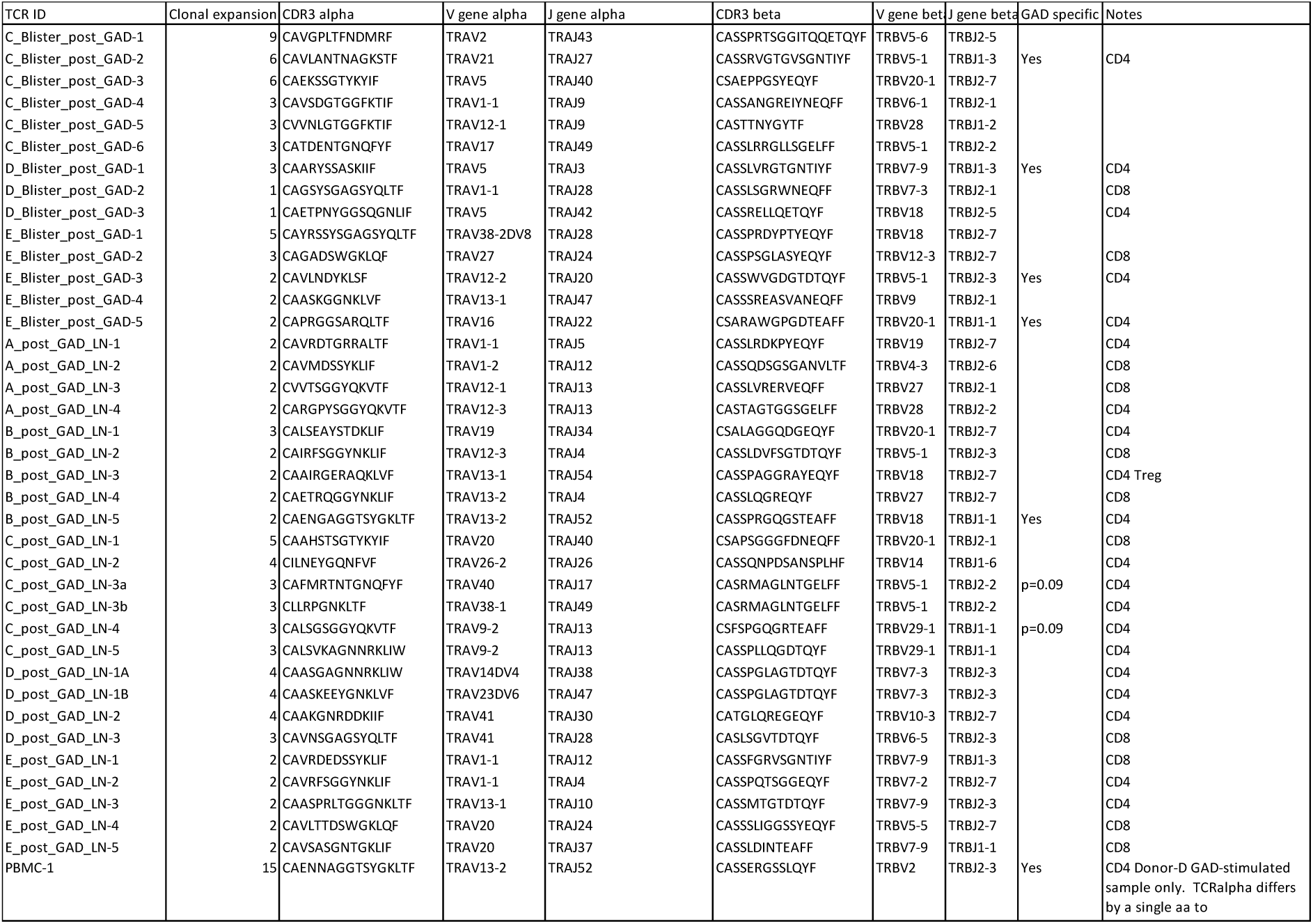
Re-expressed TCRs.

### Re-expression of lymph node TCRs identified GAD-specific T cells

From the LN clonal expansions, we re-expressed the most abundant post-injection TCRs and assessed their GAD reactivity *in vitro* using a 5KC reporter system. TCR transductants were initially screened with GAD-Alum, and those showing significant NFAT reporter response (not shown) were subsequently screened using GAD-expressing LCLs. Of the 24 TCRs re-expressed from 22 T cell clones studied (two T cell clones each expressing two TCRα chains), one clonotype (∼4% of those tested) demonstrated GAD specificity in post-injection LN samples, (Fig 2A, Table 1) and was designated TCR B_post_GAD_LN-5.

Shared clonally-expanded TCRs detected in both LN_Post and LN_Pre samples from the same donor were not GAD specific (Table 1). All confirmed GAD-specific TCRs derived from CD4+ T cells, while none of the eleven CD8+ T cell TCRs tested exhibited GAD-reactivity (Table 1). Alum alone did not induce NFAT reporter activation in 5KCs expressing these TCRs (data not shown). When mapped back onto the UMAP plot, cells expressing the same GAD-specific TCR clonotype exhibited highly similar gene expression profiles, while expression patterns differed between distinct GAD-specific clonotypes (Fig 2B).

### T cell phenotypes in the lymph node and blister

Across all samples, clonally-expanded cells were enriched within two major populations: CD8+ T cells expressing IFNG, and CD4+ T cells expressing high levels of the Th1 and Th17-associated chemokine receptors CCR6, CXCR3. We focussed particularly on Th17.1 cells, given our previous observations that reductions in this subset predict response to Ustekinumab in T1D(20). However, in the absence of *ex vivo* stimulation, these CD4+ T cells did not express IL17A (Fig 3A).

**Fig 3:**
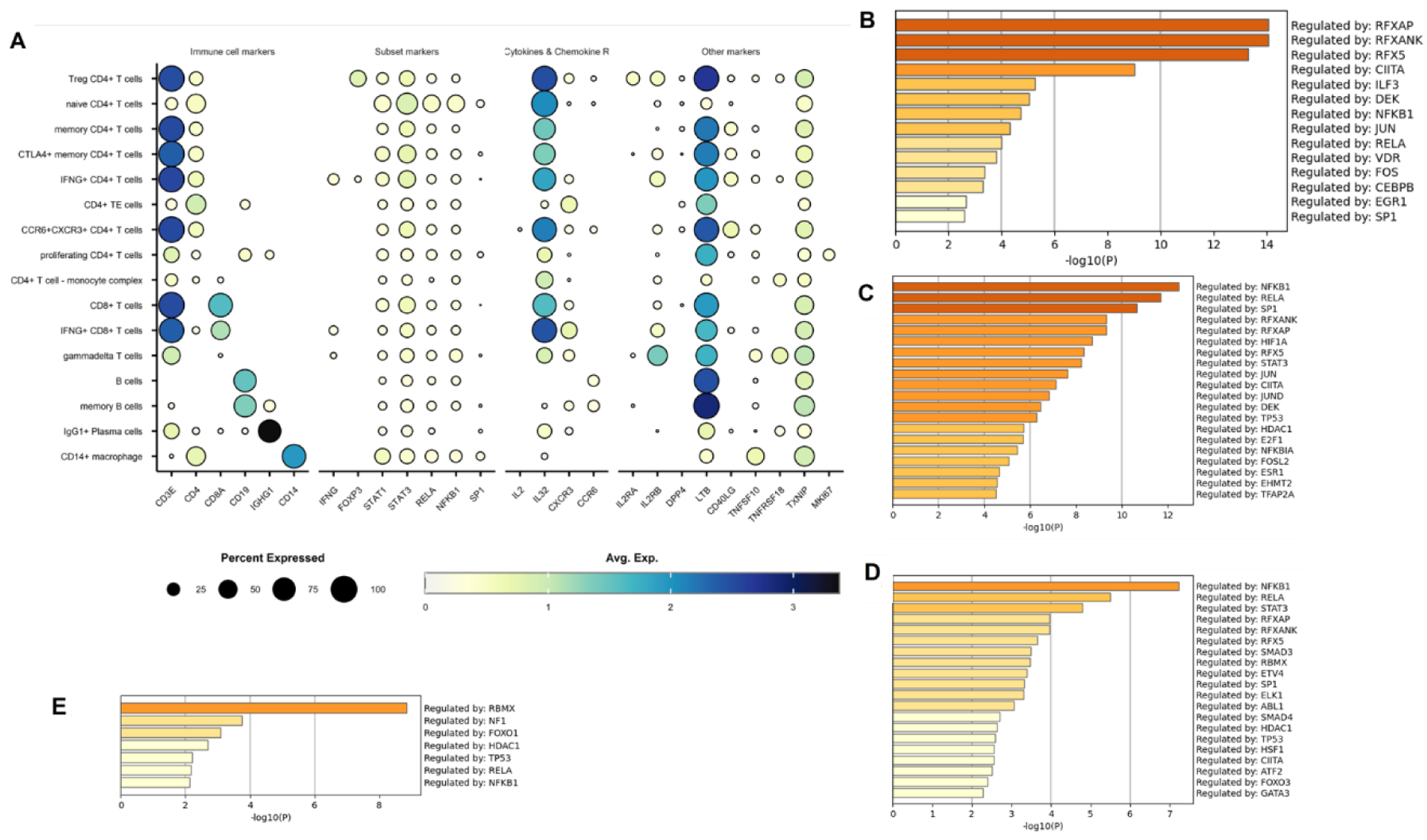
Gene expression profiles of T cells following *in vivo* GAD. **A**. DotPlot showing expression of major subset markers across scRNAseq clusters from combined LNs and blister samples **B**. Transcription factors identified by TRRUST analysis of all genes upregulated in blister T cells following GAD injection compared to control blisters **C**. Transcription factors identified by TRRUST analysis of genes upregulated in conventional CD4+ T cells from blisters compared to pre-GAD LN samples. **D**. Transcription factors identified by TRRUST analysis of all upregulated genes in clonally-expanded CD4+ T cells post-GAD vs pre-GAD **E**. Transcription factors identified by TRRUST analysis of all genes upregulated in clonally-expanded CD8+ T cells post-GAD vs pre-GAD

### Phenotypes of GAD-specific cells in the skin

Because a high proportion of the re-expressed clonally-expanded TCRs from the blisters were GAD-specific, we compared all blister T cells (with a defined TCR and not associated with antigen-presenting cells) from the GAD-injected participants to those from control donors. Following GAD injection, T-cells showed upregulation of mitochondrial and ribosomal genes, consistent with increased cellular activation. TRRUST analysis indicated that these genes were regulated by transcription factors from the RFX family (Fig 3B, Supplementary Table 5). We next examined canonical T cell subset markers of and found substantial inter-donor variability. Donor D exhibited expression of IL22, CXCR3, CCR6 and LTB; genes associated with Th17 cells and autoimmune responses(22), whereas this pattern was not observed in Donor E (Sup Fig 7).

To further characterize activation-associated changes, we compared all GAD blister T cells within conventional CD4+ T cell clusters (i.e. excluding those associated with monocytes) to CD4+ T-cells from pre-GAD LNs. The top DEGs included IL22, CXCR6 and CD26 as well as activation markers such as TNFRSF18 (GITR) (Supplementary Table 6). These changes were primarily driven by high expression in Donor D, who contributed more conventional CD4+ T cells than Donor E (Supplementary Fig 8). TRRUST analysis identified NFkB, RelA and SP1 (which regulates TBET expression(23)) as the top three transcription factors driving the observed gene expression profile in blister T cells (Fig 3D)

### Phenotypes of T cells in the lymph nodes following intradermal GAD-Alum

We next examined changes in clonally-expanded T cells following GAD-Alum injection, hypothesising that recruitment of GAD-specific T cells into the clonally-expanded T cell clusters would generate a detectable transcriptional signature. Focussing first on CD4+ clonally-expanded T cells, differential expression analysis of post-GAD samples revealed upregulation of ribosomal genes consistent with increased transcriptional activity, and of TXNIP. TRRUST analysis performed using Metascape(24) on upregulated genes (uncorrected p <0.05) identified NFkB, RelA and STAT3 as the top three transcription factors driving the post-GAD signature (Fig 2B, Supplementary Table 8). Genes regulated by these factors included TXNIP and members of the TNF superfamily. When examined by donor, TXNIP upregulation was particularly consistent across individuals (Supplementary Fig 9).

We next analysed clonally-expanded cells in the IFNG+ CD8+ T cell cluster. Again, the most significantly upregulated genes were associated with increased activation and protein translation. TRRUST analysis identified RBMX, NF1 and FOXO1 as key transcriptional regulators of this signature (Fig 2C, Supplementary Table 9).

### Comparison of gene expression in GAD-specific peripheral blood T cells reveals distinctive autoantigenic phenotypes

We next investigated whether GAD-specific cells could be identified in the peripheral blood. To this end, we performed an *in vitro* AIM assay to detect GAD-reactive CD4+ T cells prior to GAD injection. Unstimulated cells served as a negative control for unactivated cells, SEB-stimulated cells as a positive control, and buffer-stimulated control (PBMC_Ctrl) as a control for non-specific activation, or pre-activated responses. Sorted cells formed distinct clusters, including resting cells, Tregs, three populations of activated central memory (TCM) cells and two clusters co-expressing IL17A and IFNG, designated Th17.1s (Fig 4A, B). As expected, cluster frequencies differed substantially between SEB-stimulated and unstimulated conditions (data not shown). The mean stimulation index of GAD versus control stimulation, corrected for initial PBMC input, was 2.2 (range 1.2-4.9). Th17.1 clusters were enriched in IFNG, IL17A, IL17F and CSF2. Subsets designated Th17.1IL2+ and Th17.1IL22+ co-expressed IL2 or IL22, respectively (Fig 5B). These populations were of particular interest given our previous finding that reduction in Th17.1 cells predicts response to Ustekinumab in T1D(20). Compared to PBMC_Ctrl cells, PBMC_GAD cells exhibited increased expression of IFNG, IL17A, IL17F, CSF2, IL2 and IL22 as well as the Th1- and Th17-associated transcription factors STAT1 and STAT3 (Fig 3B). Pairwise comparisons between control and GAD stimulation for each donor showed this pattern of particularly elevated IL17A, IL17F and IL22 expression, was consistent across all donors except Donor B, who displayed the lowest stimulation index and an insufficient number of surviving cells after scRNAseq quality control (Supplementary Fig 10). Despite these transcriptional changes, the frequency of cells within individual clusters did not increase. Differential gene expression analysis across all cells revealed PBMC_GAD cells were defined by upregulation of IL2RA, IL2RB and LTB, DPP4 (CD26) and IL22 (Fig 3B, Supplementary Table 8). TRRUST analysis identified Rela, STAT1 and STAT3 as the top transcription factors driving the gene expression programme in GAD-reactive cells (Fig 3C).

**Figure 4.**
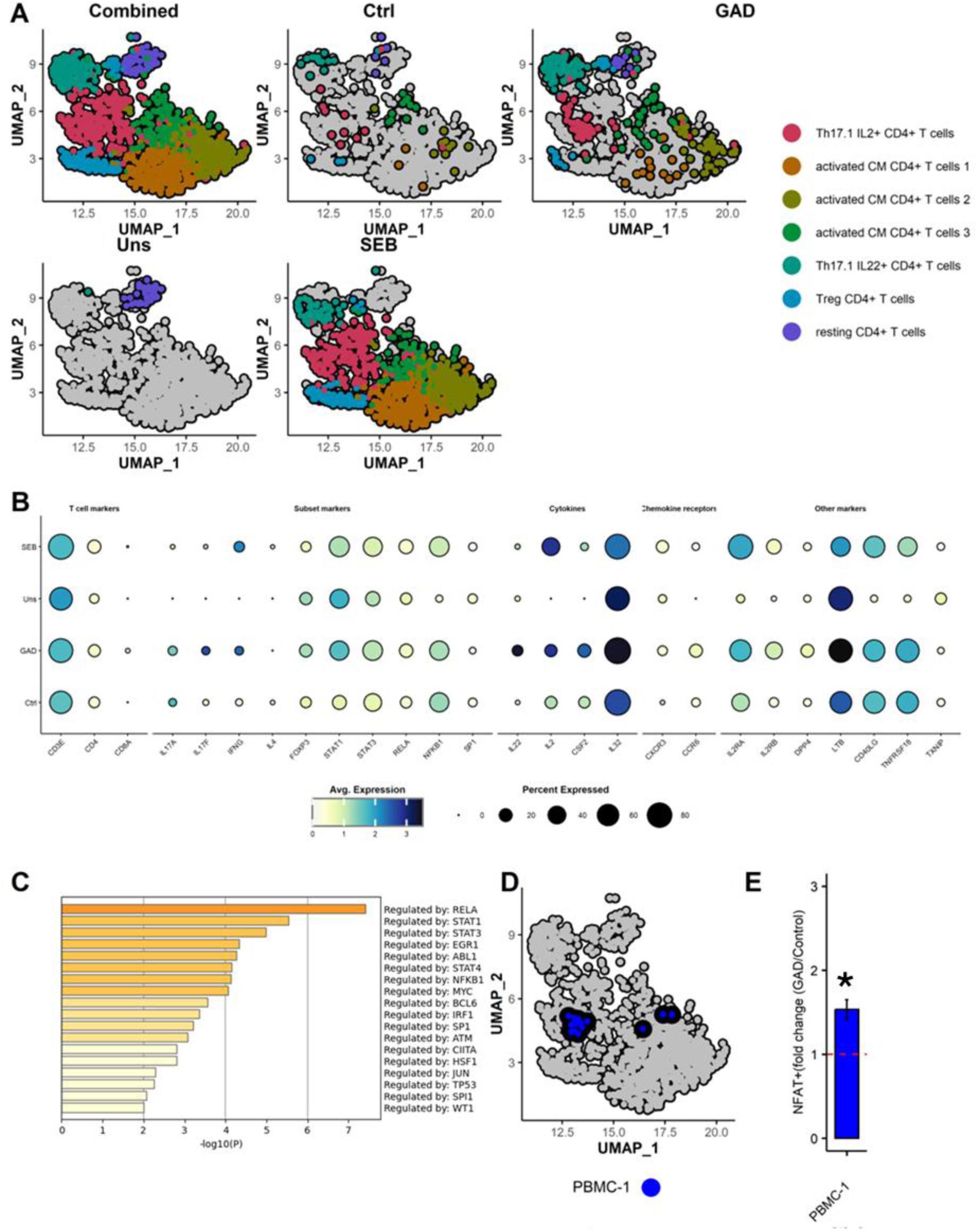
GAD-specific T cells from peripheral blood have Th17.1 phenotypes. **A**. UMAP plot of all flow-sorted cells separated by *in vitro* restimulation condition **B**. Dot plots showing expression of key markers across each cluster. **C**. Transcription factors identified by TRRUST analysis of genes upregulated in GAD-activated compared to control-activated cells. **D.** UMAP plot highlighting a clonal expanded population of cells (“PBMC-1”) detected only in the GAD-activated sample. **E**. PBMC-1 TCR re-expressed in 5-KC reporter cells and stimulated with control or GAD-expressing LCLs *p<0.05 unpaired t test.

One large clonal expansion of GAD-specific cells was detected in Donor D (15 cells, Fig 3D). These cells predominantly exhibited a Th17.1IL2+ phenotype and re-expression of their TCR in the 5KC reporter line confirmed GAD specificity (Table 1, Fig 3E). Interestingly, despite being from a different donor (Donor_B), the TCR alpha chain of the LN-derived clone B_post_GAD_LN-5 (CAEN**G**AGGTSYGKLTF) differed from that of the PBMC clone PBMC-1 (Donor_D CAEN**N**AGGTSYGKLTF) by only a single amino acid. These donors shared several HLA alleles, including HLA-DRB1*04:01:01.

Together, these findings demonstrate that GAD-specific CD4+ T cells can be isolated and characterised from peripheral blood, skin, and lymph node samples following intradermal GAD-Alum administration. Across compartments, these cells displayed transcriptional features consistent with Th1 and Th17.1 phenotypes, including activation of NFκB-, RelA-, and STAT-driven pathways. The identification of clonally-expanded, GAD-reactive TCRs in both peripheral and lymphoid compartments provides new opportunities to track autoantigen-specific T cells longitudinally and to evaluate their modulation during immunotherapeutic intervention.

## Discussion

In this study, we addressed the challenge of tracking antigen-specific cells in T1D by combining intradermal injection of diabetes autoantigen with sampling from multiple accessible sites of antigen interaction. Following intradermal GAD-Alum injection, we observed striking clonal expansion of T cells in the skin with 20-45% of all T cells clonally-expanded and up to 15 copies of individual clones. Re-expression of these TCRs confirmed that ∼30% were GAD-specific, representing approximately 1 in 10 T cells recovered from the skin injection site. In the draining LN around 2% of T cells were clonally-expanded and 4% of these were GAD-reactive, equivalent to 1 in 1000 LN T cells. Clonally-expanded LN T cells also showed transcriptional changes consistent with antigen-driven activation. In peripheral blood, clonal expansions were rare, but one clonotype was confirmed GAD-reactive. Brief *in vitro* restimulation identified GAD-reactive CD4+ T-cells representing around 0.01% of blood T cells. The identification of multiple GAD-specific TCRs offers the opportunity to track these “sentinel” clonotypes longitudinally as markers of disease activity.

Antigen-specific T cell enrichment at sites of disease is well recognised in autoimmunity(25–27), including T1D where expanded T cells in the pancreas and pancreatic LN are enriched for diabetes autoantigen specificity(28–30). Our data show that similar enrichment can be replicated in the skin following intradermal autoantigen injection, providing a minimally invasive means of monitoring disease-relevant responses. The frequency of GAD reactive CD4+ T cells compares favourably with reports from pancreatic islets where 14 of 187 TCR clonotypes responded to preproinsulin peptides(31). The relative paucity of CD8+ GAD-reactive cells was unexpected given that 3-40% of islet CD8+ T cells in T1D recognise proinsulin(30), but this may reflect donor HLA type or differences in CD4+ and CD8+ trafficking. In earlier work using a proinsulin-gold nanoparticle conjugates we detected both GNP-reactive and proinsulin reactive CD8+ T cells months after injection(18), suggesting kinetics may also be important.

This approach offers several advantages over established methods such as ELISPOTs, HLA tetramers and *in vitro* expansion assays(3, 7, 32). Unlike these techniques, scRNAseq allows full transcriptional profiling of each antigen-specific T cell without preselecting functions or markers. It does not require prior knowledge of HLA restriction or defined epitopes and, through re-expression of paired TCRαβ chains in reporter lines, enables direct confirmation of antigen specificity independent of phenotype. This design also avoids the bias introduced by repeated in vitro stimulation, which can preferentially expand highly proliferative clones and distort their native phenotype.

Across skin, LNs and blood, GAD-reactive CD4+ T cells shared a conserved transcriptional programme, dominated by NFkB, RelA, STAT1 and STAT3 transcription factors and which are associated with Th17 and Th1 differentiation(33, 34). These cells expressed CCR6 and CXCR3 consistent with Th17.1-like phenotype and the ability to migrate to islets. Interestingly, only GAD-reactive blood T-cells expressed IFNG and IL17A, likely reflecting the timing of antigen exposure to GAD (recent *in vitro* restimulation versus earlier *in vivo* activation). We also identified TXNIP upregulation in post-GAD LNs, aligning with reports of its elevation at T1D onset and modulation by verapamil.(35)

Despite these strengths, several caveats remain. The phenotypic heterogeneity in the phenotypes of GAD-specific T cells between donors and compartments may reflect differences in disease course or sampling limitations(36). In addition, the T cell clones detected peripherally may not perfectly represent those active in the pancreas, a limitation shared with all peripheral monitoring approaches. Nevertheless, the identification of IFNG+IL17A+CSF2+ CD4+ T cells in response to GAD is notable as similar cells have been linked to accelerated C-peptide loss during alefacept therapy(37) and implicated in other autoimmune diseases(38–40). We recently reported that circulating Th17.1 cells co-expressing IL-2 and GM-CSF are targeted by Ustekinumab and that reductions in this subset correlate with improved C-peptide preservation in T1D(20) This suggests that the GAD-reactive Th17.1-like cells identified here may have direct disease relevance.

GAD-reactive CD4+ T cells also expressed high levels of IL-32, previously associated with T1D progression(41) (42, 43) and IL-22, a Th17-associated cytokine elevated in human T1D(44, 45). Although IL-22 does not appear to directly drive beta-cell destruction in NOD mice(46) (47), it may contribute to tissue inflammation in human disease. We also observed nearly identical TRA sequences differing by only one amino acid among expanded GAD-reactive clonotypes from different donors. Public TCRs with such minimal junctional differences have been reported inT1D(48) (49), supporting convergent recognition of shared autoantigenic determinants.

In summary, we describe three complementary methods for enriching and monitoring diabetes autoantigen-specific cells in humans. Sampling from the skin after intradermal antigen challenge enables isolation and molecular profiling of clonally-expanded antigen-specific T cells, while lymph node aspiration and peripheral AIM assays provide complementary phenotypic data. Together, these methods provide a framework for longitudinal monitoring of antigen-specific T cells during immunotherapy and may accelerate the development of disease-modifying treatments for T1D.

## MATERIALS AND METHODS

### Sex as a biological variable

Our study examined males and females, and similar findings are reported for both sexes

### Study design

After obtaining informed consent, nine participants aged 18-50 years with T1D and positive for anti-GAD antibody were recruited. Their demographic characteristics are shown in Sup Table 1. Six participants received single intradermal administration of 4µg of GAD-Alum in 100µL of volume (clinical grade, Diamyd Medical) into the deltoid region. Ultrasound-guided fine needle aspiration (FNA) of the axillary LN was performed by an expert radiologist before, and 2-7 days after GAD-Alum administration in 5 participants. Out of 5 participants who received GAD-Alum and had FNA, 3 also had a skin suction blister induced 5-13 days after the antigen administration, in order to sample lymphocytes from the site of injection. Skin blisters were harvested at the time of erythema and induration over the injection site as described below.

In addition, three further participants who did not receive GAD-Alum challenge had a suction blister taken from the unchallenged skin of the deltoid region for the purpose of a negative control for the skin lymphocyte analysis (Sup table 2).

### Blood samples

PBMCs were isolated on Ficoll–Paque Plus (GE Healthcare Biosciences, Sweden) gradient centrifugation from the peripheral blood of participants immediately prior to injection of GAD-Alum. Samples were cryopreserved in CryoStor® (Sigma-Aldrich, Gillingham, UK) on the day of the collection and thawed on the day of the experiment. HLA typing was performed by the Anthony Nolan Trust (Sup Table 3)

### Suction blisters

Skin suction blisters were raised as described previously(18, 50). Briefly, suction cups were applied to the deltoid region of participants’ arms, at the site of previous injection. Skin suction blisters were performed by gradually applying negative pressure (up to 60 kPa) from a suction pump machine VP25 (Eschmann, Lancing, UK) through a suction blister cup with a 15-mm hole in the base (UHBristol NHS Foundation Trust Medical Engineering Department, Bristol, UK) for 2–4 hr until a unilocular blister had formed within the cup. The cup was left in place for a further 30-60 minutes to encourage migration of lymphocytes into the blister fluid. The cup was then removed, and the blister fluid aspirated using a needle and syringe. Blister fluid was immediately suspended in 10ml heparinised RPMI (Gibco)+ 5% foetal calf serum. Cells were counted then washed once and resuspended in an appropriate volume for scRNAseq. Samples were used fresh on the day of the collection.

### Ultrasound-guided Fine needle aspiration of LNs

Ultrasound-guided FNA was performed as previously reported(17). Under aseptic conditions, the skin and s.c. tissues down to the identified LN were infiltrated by 1–2 ml 1% (w/v) xylocaine with 1:200,000 adrenaline. Under real-time ultrasound visualization, using a 21-gauge needle and a 5-ml syringe, the LN cortex was sampled. One to two passages were used per sample. The LN sample was immediately transferred to tissue culture medium—5% (v/v) FCS (Biosera) in RPMI 1640 medium (Gibco) and transported to the laboratory at ambient temperature. PBMC were isolated using Lymphoprep (StemCell Technologies) and frozen in Cryostor CS10 (Sigma) until required.

### *In vitro* stimulation and flow cytometric sorting of CD4+ T cells

The Activation-Induced Marker AIM assay was performed similarly to previously published(7) (Sup Fig 1). Briefly PBMC from donors pre *in vivo* injection of GAD Alum, were thawed and stimulated for 18 hours in one of four conditions in X-VIVO media with 10% human AB serum at 4x10^6^ cells/ml(Merck): 1. Unstimulated 2. Whole GAD (Diamyd) (no Alum) in buffer (final concentration 10μg/ml). 3.Control-using the GAD dilution buffer only 4. Staphylococcal Enterotoxin B (SEB) (final concentration 50ng/ml). Following this, cells were stained with a cocktail of flow cytometry, cell hashing and CITE-seq antibodies. (Sup Table 4). CD4+ T cells were then sorted as follows (Sup Fig 2): SEB, GAD and buffer control-Effector T cells: CD154+ and either CD69+ or CD137+ or both; Tregs: CD154- CD137+ and either CD69+ or GARP+ or both. For GAD and buffer control conditions - all cells were sorted from the positive gate. For SEB-stimulated - 100 activated Tregs and 1000 activated effectors were sorted. From the unstimulated condition, 1000 cells negative for all four activation markers were sorted. All cells were processed for scRNAseq on the day of sorting.

### scRNAseq

LN and injected blister samples were processed using a 10xGenomics V1 human 5’GEX kit and libraries were constructed following standard 10x Genomics protocols. Samples were pooled and sequenced on Illumina NextSeq and MiSeq. Demultiplexed samples were processed individually using the 10xGenomics CellRanger 3.0.2 pipeline and GRCh38-1.2.0 with vdj GrCh38-alts-ensembl 2.0.0. For control blisters and PBMCs the 10xGenomics V2 human 5’GEX kit and CellRanger 6.1.1 pipeline were used. Data were further processed using the Seurat V4 package in R(51) and DoubletFinder(52) and manual gating were used to remove doublets. TCR sequences and clone numbers were added to the Seurat object metadata slot. TCR clone numbers refer to data generated in the CellRanger pipeline, even if clones of the same cell did not meet gene expression quality control thresholds. Raw data contained red blood cells, but these were removed automatically in the QC process due to low levels of UMIs and low unique genes identified. Cell clusters were determined by Seurat. We performed batch correction with Harmony(53) to produce a single Seurat object. Data were further analysed in R using scPubr, ggplot2, cowplot, ggpubr and tidyverse packages. Differentially expressed genes (DEGs) were identified in Seurat and TRRUST pathways were analysed in Metascape v3.5.20240901.

### Antigen specificity experiments

TCR specificity was determined as previously described(8, 18). Briefly α-Chain and β-chain genes connected by the porcine teschovirus-1 2A peptide gene were engineered into a murine stem cell virus–based retroviral vector. T-cell transductants were generated by retroviral transduction of the 5KC murine T-cell hybridoma cells lacking endogenous TCR expression, expressing fluorescent proteins downstream of the NFAT promoter and expressing human CD4 or CD8 co-receptors according to the original T cell phenotype. 5KC cells were spin-infected with replication-incompetent retroviruses that were produced from Phoenix cells transfected with the retroviral vectors and the pCL-Eco packaging vector. Cells transduced with retroviruses were then selected by sorting for CD3 expression. (CD3ε MicroBead Kit, mouse, Miltenyi Biotec).

Codon optimised *GAD2* gene (GAD/GAD65 protein) was cloned into the third-generation lentiviral transfer vector backbone pELNS (kindly provided by Dr. James Riley, University of Pennsylvania, PA, USA) containing a rat CD2 (rCD2) marker gene preceded by a P2A self-cleaving sequence: XbaI-Kozak-*GAD2*-XhoI-P2A-rCD2-SalI-Stop. Lentiviral particles were produced using HEK293T cells and envelope plasmid pMD2.G (gift from Didier Trono, Addgene plasmid #12259), and packaging plasmids pMDLg/pRRE (gift from Didier Trono, Addgene plasmid #12251) and pRSV-Rev (gift from Didier Trono, Addgene plasmid #12253) and used to insert GAD into participants’ EBV-transformed B cells (lymphoblastoid cell lines, LCL) 5KC cells bearing TCRs were incubated for 48h with autologous LCLs with antigens as indicated at 1x10^6^ 5KC/ml and 1x10^6^ LCLs/ml in 200µL RPMI 1640 (Gibco)+penicillin/streptomycin (Gibco) +10% foetal calf serum (BioSera)+ 2-Mercaptoethanol (Thermo Fisher Scientific) in 96 well round bottomed plates. The positive control was 5µL αCD3/CD28 microbeads (Dynabead). Flow cytometry was performed on a BD Canto II. Flow cytometry data was analysed using FlowJo 10.7.1. Murine IL-2 ELISAs were performed on undiluted cell culture media following manufacturer’s instructions (Biolegend).

### Statistics

DEGs that defined cell clusters in Seurat were determined using Seurat’s inbuilt statistical methods using the Wilcoxon rank sum test, using either only.pos=TRUE or only.pos=FALSE as described in the results section. DEG pathway analysis was conducted using Metascape standard parameters(54) on all DEG with an uncorrected p< 0.05. TCR-5KC reactivity to GAD-Alum presented by EBV transformed autologous B cells (LCLs), or GAD expressing LCLs was determined using a one tailed T test compared to TCR-5KC with LCLs alone.

### Study approval

This study was carried out with the approval of the UK Research Ethics Service (Wales Research Ethics Committee 1, reference number 13/WA/0113). Written informed consent was obtained from all participants. The study was conducted in compliance with the principles of the Declaration of Helsinki (1996) and the principles of Good Clinical Practice and in accordance with all applicable regulatory requirements including but not limited to the Research Governance Framework and the Medicines for Human Use (Clinical Trial) Regulations 2004, as amended in 2006. Written informed consent was received for the use of photographs and the record of informed consent has been retained.

## Supporting information

Supporting Data values

## Data availability

scRNAseq data will be deposited in a publicly available repository at the point of publication.

## Author contributions

SJH: Conceptualization, Data curation, Formal Analysis, Funding acquisition, Investigation, Methodology, Visualization, Writing – original draft, Writing – review & editing; EJSR: Investigation, Methodology, Writing – review & editing; TCT: Investigation, Methodology, Writing – review & editing; MN: Methodology, Resources, Writing – review & editing; LL: Methodology, Resources, Writing – review & editing; RA: Data curation, Software, Writing – review & editing; GD: Methodology, Resources, Writing – review & editing; JD: Investigation, Project administration, Writing – review & editing; EW: Investigation, Methodology, Writing – review & editing; JAP: Funding acquisition, Methodology, Supervision, Writing – review & editing; AKS: Methodology, Resources, Writing – review & editing; PN: Methodology, Writing – review & editing; DW: Methodology, Writing – review & editing; AH: Investigation, Project administration, Writing – review & editing; PY: Investigation, Methodology, Writing – review & editing; MH: Investigation, Methodology, Writing – review & editing; AL: Methodology, Resources, Writing – review & editing; FSW: Supervision, Writing – review & editing; TIMT: Investigation, Methodology, Resources, Writing – review & editing; CMD: Conceptualization, Funding acquisition, Supervision, Writing – review & editing; DT: Conceptualization, Funding acquisition, Writing – review & editing

## Acknowledgments

We thank Dr. James Riley, University of Pennsylvania, PA, USA for the gift of the third-generation lentiviral transfer vector backbone, pELNS.

