## Supplemental Figures for "Tracking GAD-specific T-cell expansions in Type 1 diabetes by intradermal GAD-Alum challenge"

Supplementary figures

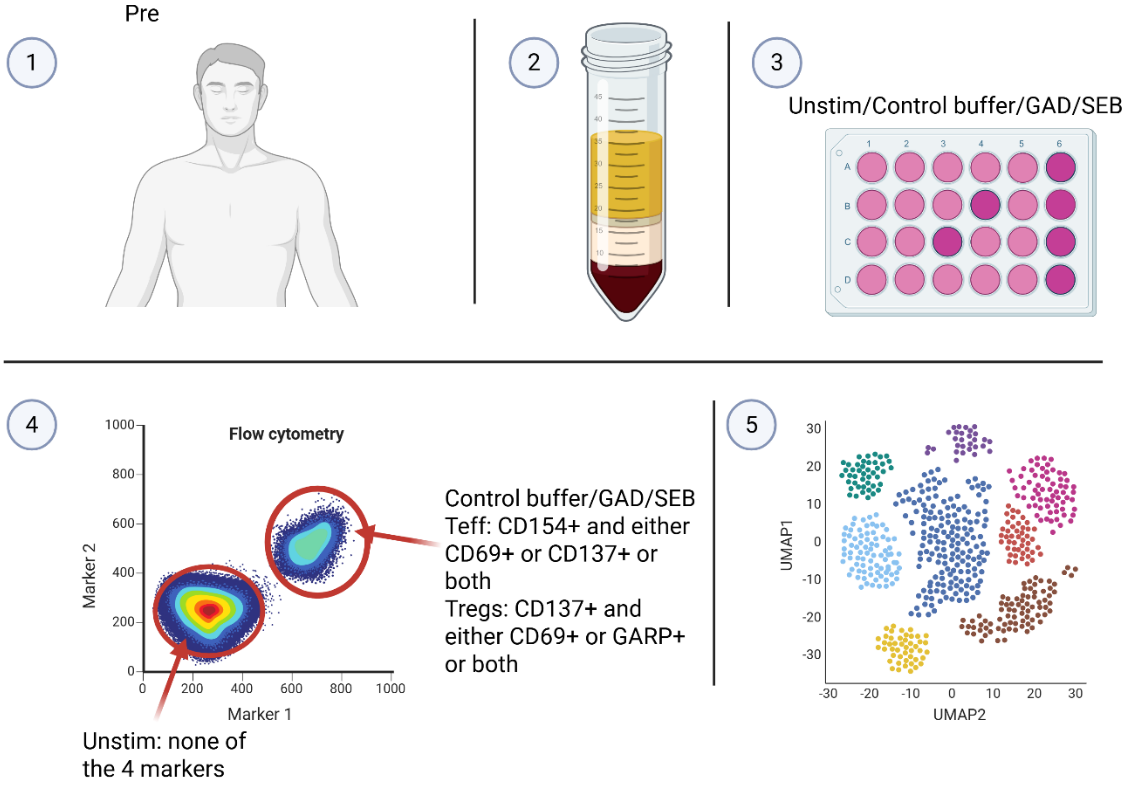


**SupplementaryFigure 1: Outline of the AIM assay**

**1**. Participants were sampled before (**Pre**) *in vivo* injection of GAD-Alum. **2**. Peripheral blood mononuclear cells (PBMCs) were isolated and cryopreserved until required. **3**. PBMCs were thawed and cultured for 18h under one of four conditions. *Unstim*- unstimulated cells cultured in media alone. *Control Buffer* – cells cultured with buffer only. *GAD* – cells cultured with GAD in buffer. *SEB*- cells cultured with the superantigen SEB. **4**. From the Unstim wells, unactivated CD4+ T cells were sorted (neagative for all four activation markers). From Control buffer/GAD and SEB wells, activated CD4+ effector T cells (CD154+ and either CD69+ or CD137+ or both) and activated CD4+Tregs (CD137+ and either CD69+ or GARP+ or both) were isolated. The detailed flow cytometry gating strategy is shown in Supplementary Figure 2. **5**. Sorted cells were subjected to scRNAseq.

Created in BioRender. H, S. (2026) https://BioRender.com/45z5gll

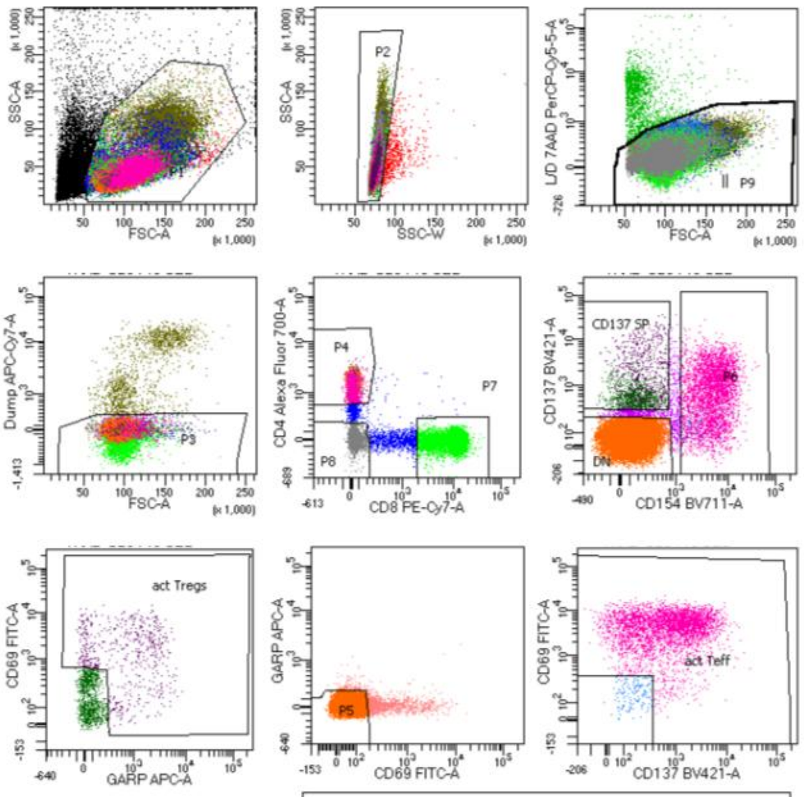


| Population | #Events | %Parent | %Total |
|---|---|---|---|
| All Events | 123,841 | #### | 100.0 |
| P1 | 109,158 | 88.1 | 88.1 |
| P2 | 108,490 | 99.4 | 87.6 |
| P9 | 106,521 | 98.2 | 86.0 |
| P3 | 91,192 | 85.6 | 73.6 |
| P4 | 40,080 | 44.0 | 32.4 |
| DN | 33,564 | 83.7 | 27.1 |
| P5 | 32,018 | 95.4 | 25.9 |
| CD137 SP | 1,584 | 4.0 | 1.3 |
| act Tregs | 609 | 38.4 | 0.5 |
| P6 | 4,037 | 10.1 | 3.3 |
| act Teff | 3,896 | 96.5 | 3.1 |
| P7 | 32,823 | 36.0 | 26.5 |
| P8 | 10,875 | 11.9 | 8.8 |

**Supplementary Figure 2.** Flow cytometry gating strategy (SEB-stimulated example). Representative gating strategy used to identify activated CD4+ T-cell subsets following SEB stimulation. Shown are sequential gates for live singlet lymphocytes, CD3+CD4+ T cells, and activated effector or regulatory populations based on expression of CD154, CD69, CD137 and GARP. This gating strategy was applied consistently to all stimulation conditions (Unstim, Control buffer, GAD and SEB) to define activated CD4+ Teff and Treg subsets for subsequent single-cell RNA sequencing.

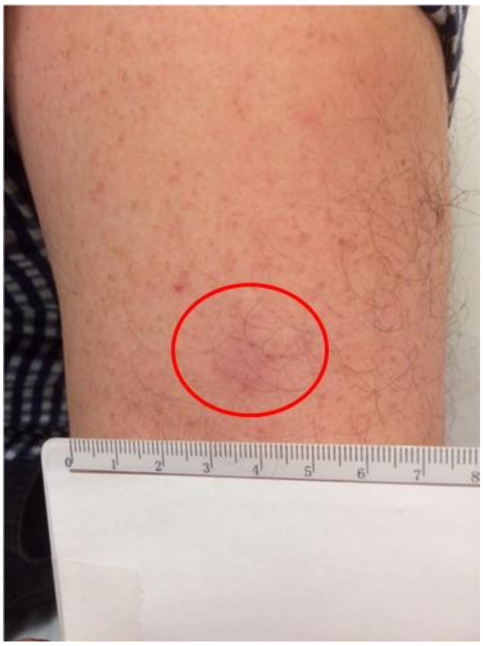


**Supplementary Figure 3. Example of delayed skin reaction following intradermal GAD-Alum injection.**
Representative clinical photograph showing the characteristic delayed erythematous induration (2–3 cm diameter) that developed at the site of intradermal GAD-Alum injection. The reaction typically appeared 2–3 days post-injection and resolved spontaneously within 15 days.

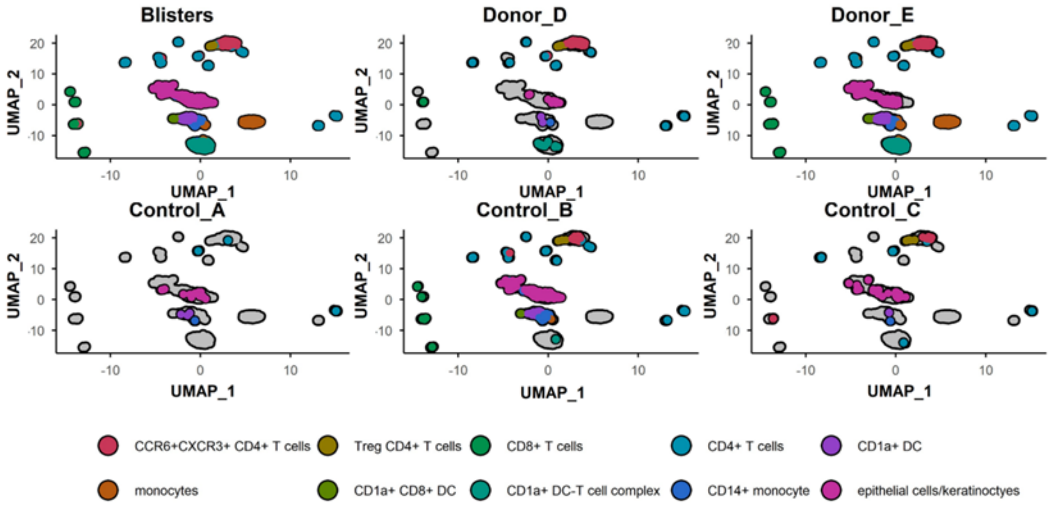


**Supplementary Figure 4. Single-cell RNA sequencing of cells from GAD-injected and control blisters.** UMAP plots of all cells from GAD-injected and control blisters, clustered in Seuratand coloured by cell type. Data are shown both combined and separated by donor.

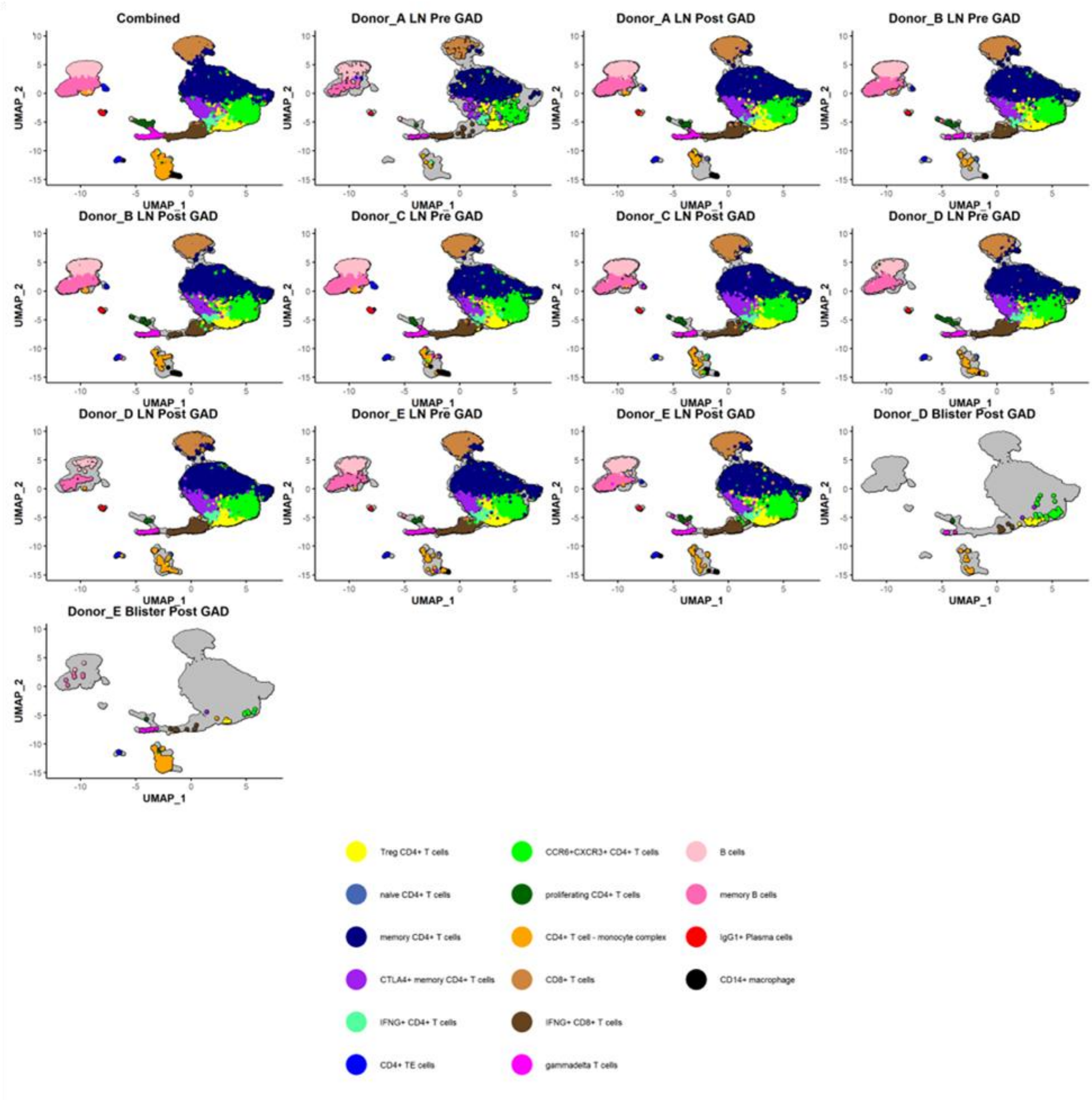


**Supplementary Figure 5. Single-cell RNA sequencing of cells from GAD-injected blister and pre- and post-GAD lymph nodes.**

UMAP plots of all cells from GAD-injected blisters and lymph node samples collected before and after GAD injection. Cells were clustered in Seurat and coloured by cluster identity. Combined data are shown, followed by plots split by donor and sampling timepoint.

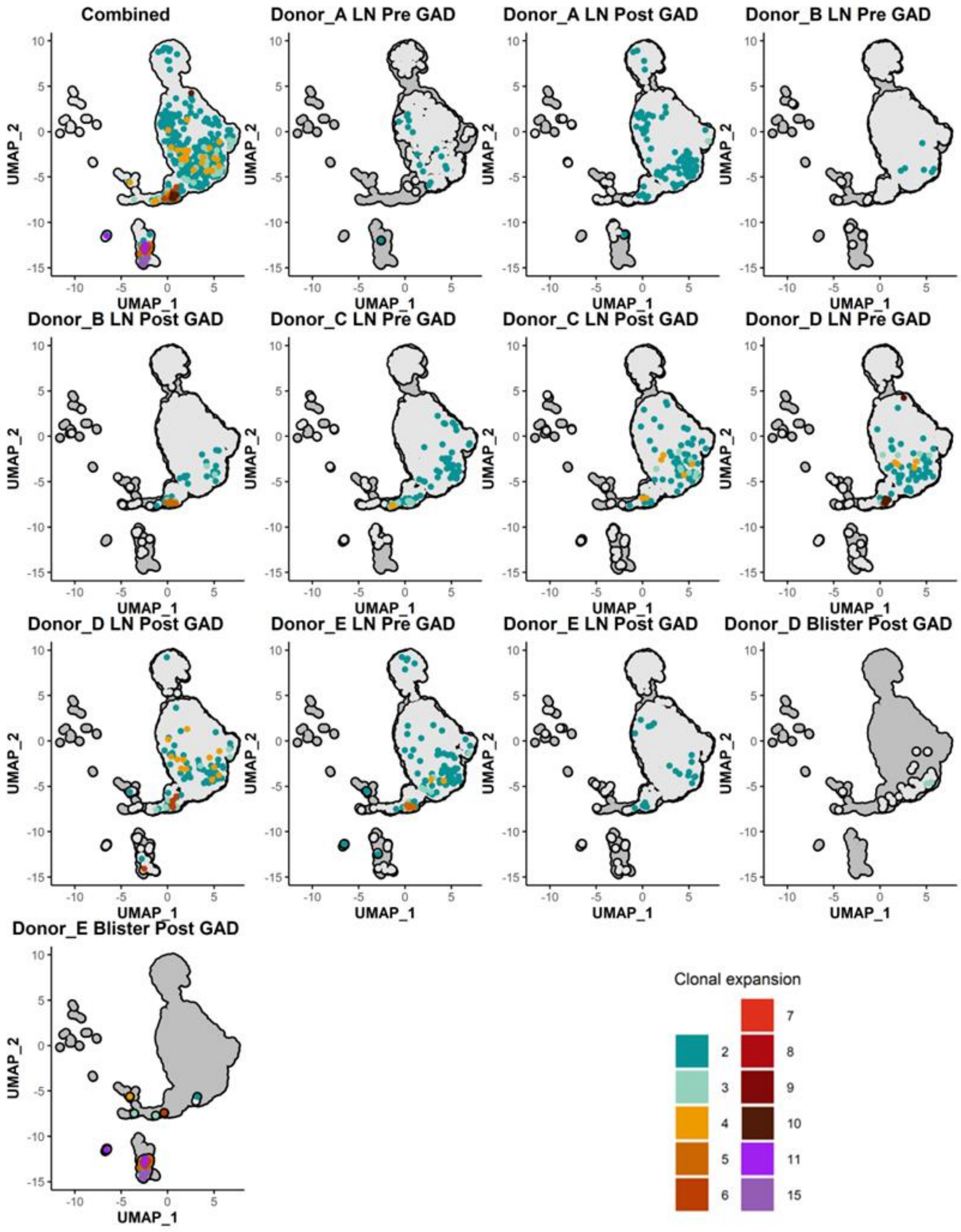


**Supplementary Figure 6. Clonal expansions of TCRs in lymph nodes and blisters.** UMAP plots of all cells from lymph node and blister samples, clustered in Seurat. Combined data are shown, followed by plots split by donor and sampling timepoint. Cells are coloured according to TCR clonal expansion.

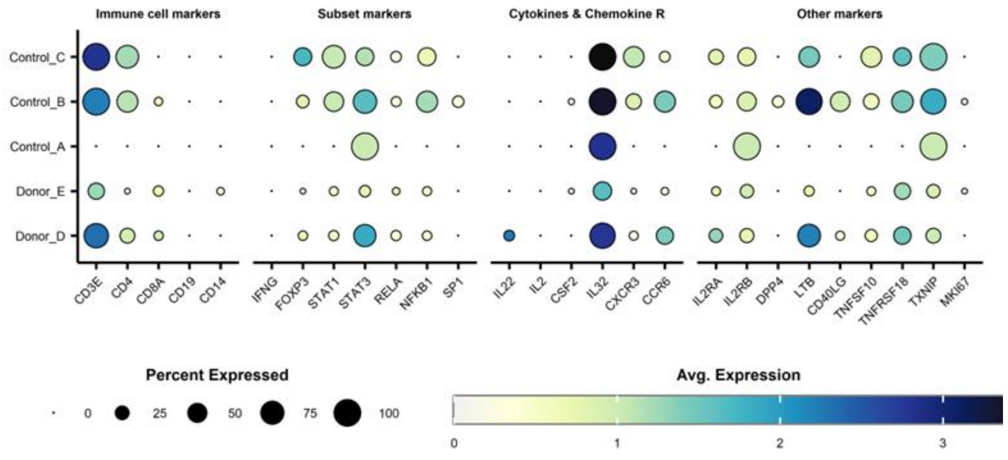


**Supplementary Figure 7. Comparison of gene expression in CD4+ T cells from blisters.**

Gene expression profiles of CD4+ T cells from blister samples, shown separately for each donor to illustrate inter-individual variation in transcriptional signatures.

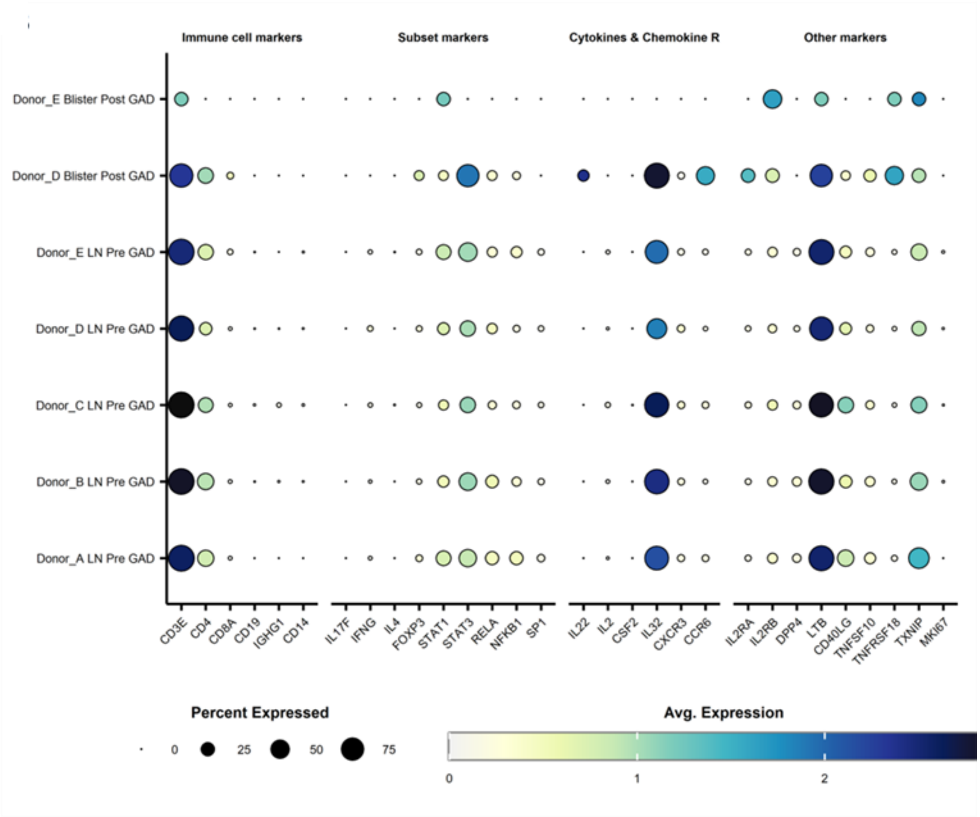


**Supplementary Figure 8. Comparison of gene expression in CD4+ T cells from blisters versus pre-GAD lymph nodes.**

Differential gene expression profiles of CD4+ T cells from GAD-injected blister samples compared with pre-GAD lymph nodes, shown separately for each donor.

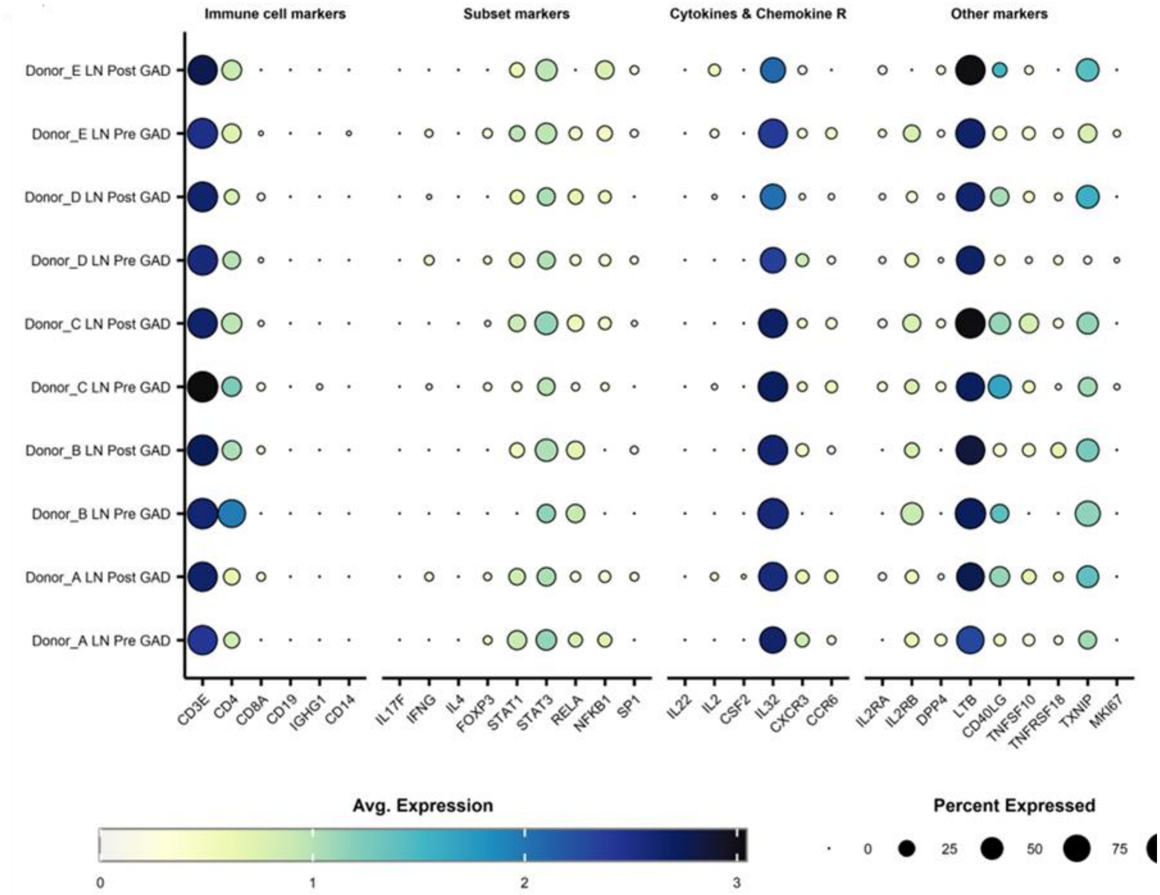


**Supplementary Figure 9. Gene expression in clonally expanded CD4+ T cells in lymph nodes.**

Gene expression profiles of clonally expanded CD4+ T cells from lymph nodes collected before and after GAD injection, highlighting transcriptional changes associated with antigen-driven activation.

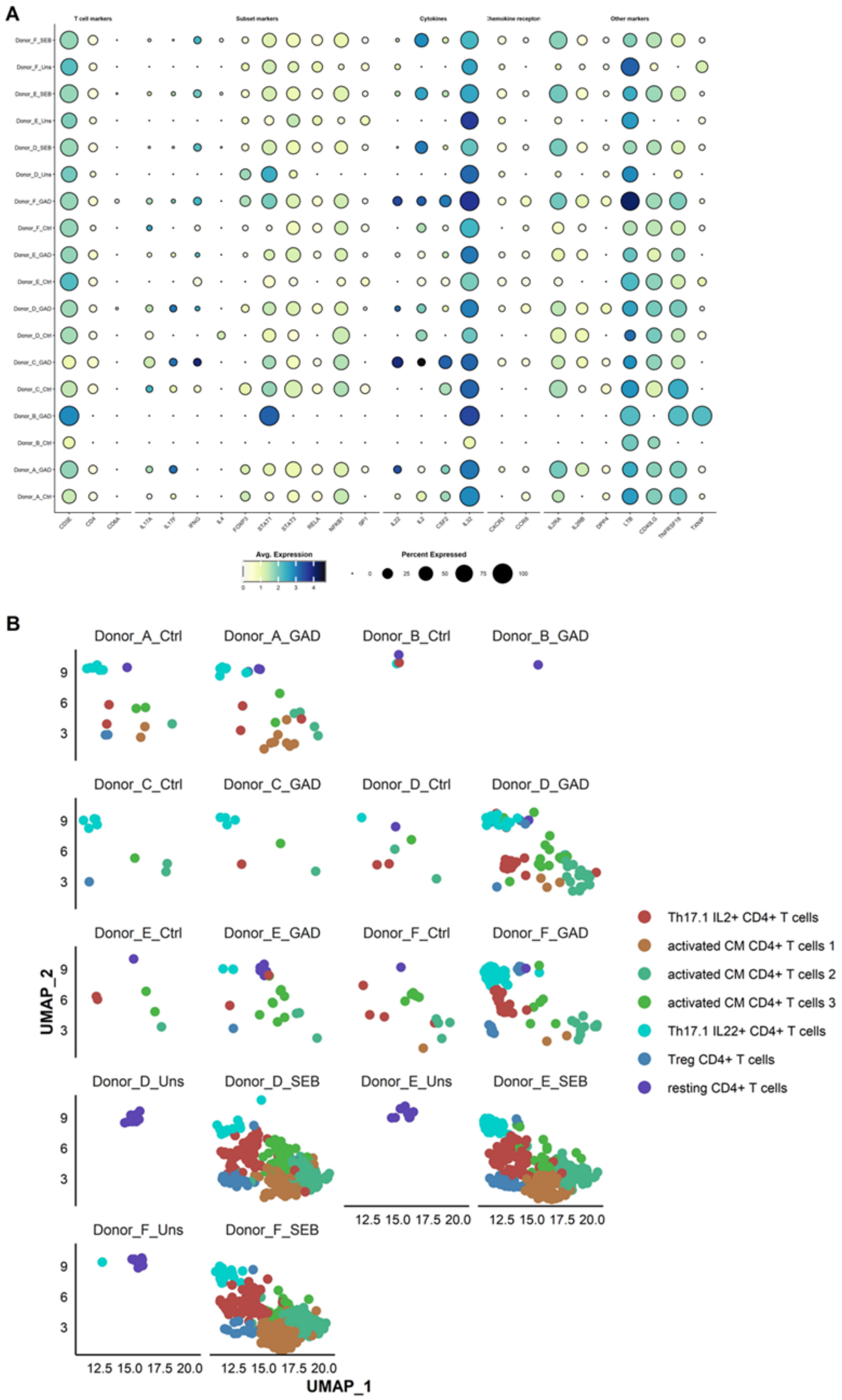


**Supplementary Figure 10. Gene expression and clustering of PBMCs following in vitro stimulation.**

**A**. Dot plot showing gene expression profiles of PBMCs, split by donor and stimulation condition. **B**. UMAP plots of all cells, split by donor and in vitro stimulation condition, illustrating clustering patterns across experimental groups.
